# Inhibition of Invasive Salmonella by Orally Administered IgA and IgG Monoclonal Antibodies

**DOI:** 10.1101/780148

**Authors:** Angelene F. Richards, Jennifer E. Doering, Shannon A. Lozito, John J. Varrone, Michael Pauly, Kevin Whaley, Larry Zeitlin, Nicholas J. Mantis

## Abstract

**Background:** Non-typhoidal *Salmonella enterica* strains, including serovar Typhimurium (STm), are an emerging cause of invasive disease among children and the immunocompromised, especially in regions of sub-Saharan Africa. STm invades the intestinal mucosa through Peyer’s patch tissues before disseminating systemically. While vaccine development efforts are ongoing, the emergence of multidrug resistant strains of STm affirms the need to seek alternative strategies to protect high-risk individuals from infection.

**Methodology/Principal Findings:** In this report we investigated the potential of an orally administered O5 serotype-specific IgA monoclonal antibody (mAb), called Sal4, to protect mice against invasive *Salmonella enterica* serovar Typhimurium (STm) infection. Sal4 IgA was delivered to mice prior to or concurrently with STm challenge. Infectivity was measured as bacterial burden in Peyer’s patch tissues one day after challenge. Using this model, we defined the minimal amount of Sal4 IgA required to significantly reduce STm uptake into Peyer’s patches. The relative efficacy of Sal4 in dimeric and secretory IgA (SIgA) forms was compared, as was a second lower avidity O5-specific IgA mAb that we produced from STm immunized mice. To assess the role of isotype in oral passive immunization, we engineered a recombinant IgG1 mAb carrying the Sal4 variable regions and evaluated its ability to block invasion of STm into epithelial cells *in vitro* and Peyer’s patch tissues.

**Conclusions/Significance:** Our results demonstrate the potential of orally administered monoclonal IgA and SIgA, but not IgG, to passively immunize against invasive Salmonella. Nonethless, the prophylactic window of IgA/SIgA in the mouse was on the order of minutes, underscoring the need to develop formulations to protect mAbs in the gastric environment and to permit sustained release in the small intestine.

**Author Summary:** The bacterium *Salmonella enterica* is responsible for significant morbidity and mortality in the developed and developing worlds. While the pathogen is most renowed as the causative agent of typhoid fever, the emergence of invasive non-typhoid strains like *S. enterica* serovar Typhimurium (STm) are of great concern because of their propensity to cause severe disease in children under the age of five. In this report, we demonstrate in a mouse model that oral administration of a monoclonal antibody targeting the surface of STm is able to prevent the bacterium from infecting gastrointestinal tissues, the first step in the dissemination process. We show that IgA antibodies (which are normally found in the gut) were far superior than an equivalent IgG antibody (normally found in blood) at protecting the intestine from STm infection. These results lay the foundation for future studies aimed at the development of an orally administered antibody cocktail capable of providing temporary immunity to pathogens like *S. enterica*.

## Introduction

Enteric bacterial pathogens constitute a major burden on global health, especially in children younger than five years of age [1, 2]. The Global Enteric Multicenter Study (GEMS) surveyed children ages 0-59 months in seven countries in sub-Saharan Africa and south Asia and identified the leading etiological agents of moderate-to-severe (MSD) and less-severe diarrhea (LSD) in this age group [1, 3]. Included on the list were *Shigella* species, *Campylobacter jejuni, Vibrio cholerae* and enterotoxigenic *E. coli* (ETEC) among others. Episodes of MSD and LSD can each have long term impacts on child health, most notably linear growth faltering. Other pathogens like multidrug resistant typhoid and invasive non-typhoidal Salmonella (iNTS) caused by different serovars of *Salmonella enterica* are also a source of infections in sub-Saharan Africa that can have short- and long-term health consequences [4–6]. The dire need for vaccines against enteric bacterial pathogens like *C. jejuni, Shigella*, ETEC and *Salmonella* has been recognized for decades, especially within military settings [2]. While there are numerous candidate vaccines under evaluation, the path forward remains challenging and alternative approaches need to be considered to combat *C. jejuni, Shigella*, ETEC and *Salmonella* in the immediate future.

With the advent of affordable and scalable platforms for the production of pathogen-specific IgG and secretory IgA (SIgA), the notion of oral passive immunization with polyclonal or monoclonal antibodies (mAbs) as a strategy to blunt diarrheal diseases in high-risk populations is gaining attention. For example, it was reported that ingestion of polyclonal hyperimmune bovine colostrum (HBC), marketed as Travelan^®^, reduces experimental traveler’s associated ETEC infection [7]. Sears and colleagues recently presented evidence that IgG and possibly IgA antibodies in Travelan^®^ and a related HBC product (IMM-24E) may exert their protective effects through arresting ETEC motility and complement-mediated killing [8]. More recently, Guintini and colleagues demonstrated that oral administration of IgG or IgA mAbs targeting a single adhesin (CfaE) were able to reduce ETEC colonization by several orders of magnitude in a mouse model [9]. In the case of invasive *Salmonella enterica* serovar Typhimurium (STm), Corthésy and colleagues reported that polyreactive secretory-like IgA/IgM mixtures were capable of reducing bacterial entry into Peyer’s patch tissues [10, 11].

While these studies represent a proof of concept that oral immunoglobulins can abrogate Salmonella infection, the amount of IgA/IgM required to achieve a reduction in bacterial burden was excessive (*i.e.*, ~10 mg of SIgA/IgM; ~ 500 mg/kg) and likely impractical if translated to a human setting. For that reason, we sought to investigate the potential benefit of a mAb-based passive immunization approach in blocking invasive *Salmonella*.

Sal4 is a well-characterized, dimeric IgA mAb originally isolated from a panel of B cell hybridomas derived from Peyer’s patch tissues of mice that had been immunized with an attenuated strain of STm [12]. Sal4 recognizes the O5-antigen of STm lipopolysaccharide (LPS) [13]. The O-antigen of STm is a tetrasaccharide consisting of galactose, rhamnose, and mannose, with an abequose (3,6 dideoy-galactose) moiety on the mannose side chain. The O5 antigen is conferred when the abequose residue is acetylated, while the O4 antigen is defined by the absence of acetylation modification [14]. Both STm O4 and O5 serotypes are invasive in mouse models of intragastric and parenteral challenge, although the actual lethal dose values may vary slightly [15].

In the so-called backpack tumor model (described elsewhere in this manuscript), it was shown that Sal4 IgA, when actively transported into the intestinal lumen of mice in form of secretory IgA (SIgA), was able to reduce STm uptake into Peyer’s patch tissues [15]. Peyer’s patches represent the point of entry for invasive strains of *Salmonella enteríca* and the bottleneck for systemic dissemination [16]. Sal4 IgA’s protective capacity was limited to the gut, as even high levels of Sal4 IgA in circulation were unable to curtail STm systemic infection in the face of a parenteral bacterial challenge [15]. Thus, Sal4 IgA limits STm infection exclusively in the context of the gastrointestinal lumen. Although the exact mechanisms by which Sal4 IgA prevents bacterial uptake into Peyer’s patch tissues have not been fully resolved, Sal4 IgA strongly promotes bacterial agglutination *in vitro* and is a potent inhibitor of STm flagella-based motility in liquid and viscous media [17].

In this report, Sal4 IgA was chosen as a prototype to investigate the potential of orally administered mAbs to passively immunize against invasive Salmonella. We first established a robust mouse model of bacterial entry into Peyer patch tissues and then used the model to compare protection afforded by Sal4 as dimeric IgA, secretory IgA and even IgG. We also generated and characterized an second O5-specific IgA mAb and compared it to Sal4 IgA *in vitro* and *in vivo*.

## Methods

### Bacterial strains and growth conditions

*Salmonella enterica* serovar Typhimurium (STm) strains used in this study are described in **Table 1**. STm ATCC 14028 was purchased from the American Type Culture Collection (Manassas, VA) [18]. S. Typhimurium strains AR04 (*zjg8101::kan oafA126::*Tn*10d-Tc fkpA-lacZ*) and AR05 (*zjg8101::kan*) are derivatives of ATCC 14028, as described [17, 19]. Unless otherwise stated, single colonies were used to inoculate sterile Luria-Bertani (LB) broth and incubated overnight at 37°C with aeration, then subcultured in fresh LB to mid-log phase (OD_600_ 0.40) before use.

**Table 1.** ST strains and derivatives used in this study.

| Strain | O-Ag | genotype | X-Gal | Reference |
| --- | --- | --- | --- | --- |
| ATCC14028 | O5 | wild type | W | [18] |
| AR05 | O5 | *zjg8101::kan* | W | [19] |
| AR04 | O4 | *zjg8101::kan oafA126::Tn10d-Tc fkpA-lacZ* | B | [19] |
| CS022 | O5 | *pho-24* (PhoP constitutive) | W | [20] |
| SJF59 | O4 | *phoP-24 oafA126:: Tn10d-Tc* | W | This study |

### Monoclonal antibodies (mAbs) and hybridomas

Antibodies used in this study are listed in **Table 2**. The B cell hybridoma cell lines secreting the monoclonal polymeric IgA antibody, Sal4, specific for O5-antigen and 2D6 IgA, specific for *V. cholerae* Ogawa LPS, were originally obtained from Dr. Marian Neutra (Children’s Hospital Boston) [15]. Purified dimeric Sal4 IgA (dIgA) and recombinant human secretory component (rSC) were associated for 1 h at room temperature to generate Sal4 SIgA, as described [21]. Chimeric Sal4 IgG1 and PB10 IgG1, specific for ricin toxin, were provided by Mapp Biopharmaceutical (San Diego, CA). 2D6 IgA and PB10 IgG1 mAbs were used as IgA and IgG1 isotype controls respectively throughout the study. The PeA3 murine B cell hybridoma secreting a monoclonal IgA against the STm O5 antigen was generated from the Peyer’s patches of BALB/c mice repeatedly immunized orally with STm, essentially as described [15]. The resulting hybridomas were screened by ELISA for reactivity with STm whole cells and purified LPS (Sigma-Aldrich, St. Louis, MO).

**Table 2.** Anti-ST and control mAbs used in this study.

| Antibody | species | form | source | epitope |
| --- | --- | --- | --- | --- |
| Sal4 | mouse | IgA | hybridoma | STm O5 Ag |
| Sal4 | mouse | SIgA | hybridoma | STm O5 Ag |
| Sal4 | chimeric | IgG1 | Nicotiana | STm O5 Ag |
| PeA3 | mouse | IgA | hybridoma | STm O5 Ag |
| 2D6 | mouse | IgA | hybridoma | *V.cholerae* LPS |
| 2D6 | chimeric | IgG | Nicotiana | *V.cholerae* LPS |

### Animal care and ethics statement

The mouse experiments described in this study were reviewed and approved by the Wadsworth Center’s Institutional Animal Care and Use Committee (IACUC) under protocol #17-428. The Wadsworth Center complies with the Public Health Service Policy on Humane Care and Use of Laboratory Animals and was issued assurance number A3183-01. The Wadsworth Center is fully accredited by the Association for Assessment and Accreditation of Laboratory Animal Care (AAALAC). Obtaining this voluntary accreditation status reflects that Wadsworth Center’s Animal Care and Use Program meets all standards required by law and goes beyond the standards as it strives to achieve excellence in animal care and use. Mice were euthanized by carbon dioxide asphyxiation followed by cervical dislocation, as recommended by the Office of Laboratory Animal Welfare (OLAW), National Institutes of Health.

### Mice

Female BALB/c mice aged 8-12 weeks were obtained from Taconic Biosciences (Rensselaer, NY) and cared for by the Wadsworth Center Animal Core Facility. All experiments were performed in strict accordance with protocols approved by the Wadsworth Center’s IACUC, as noted above.

### Enzyme-linked immunosorbent assays (ELISAs)

For direct ELISAs, 96-well NUNC MaxiSorp plate (ThermoScientific, Waltham, MA) were coated with 0.1 ml of STm LPS (1 μg/ml in sterile PBS) overnight at 4°C. Wells were blocked with PBS containing 0.1% Tween-20 (PBST) and 2% goat serum for 2 h at room temperature before washing with PBST. Plates were developed using goat anti-mouse and goat anti-human HRP-conjugated secondary IgG antibodies (final concentration of 0.5 μg/ml) and SureBlue TMB Microwell Peroxidase Substrate (KPL, Gaithersburg, MD). For the whole-bacteria ELISA, 96-well NUNC MaxiSorp plates were coated with poly-L-lysine (10 μg/ml) overnight at 4°C. Overnight cultures of STm were washed twice with PBS and then placed into each well of the microtiter plate. The plates were centrifuged two times at 500 x *g* for 3 min (rotating 180° for the second spin), and then fixed with 2% paraformaldehyde (PFA) in PBS. The bacteria-coated plate was then treated with sterile glycine (0.1 M) to quench residual PFA and ELISAs were performed as described above. All plates were read by spectrophotometry (A450) within 30 minutes of developing using a VersaMax microplate reader and SoftMaxPro 5.2 software.

### Soft agar motility assays

For the soft agar motility assays, LB medium with 0.3% Bacto™ agar (Becton Dickinson) was prepared with 15 μg/ml of each mAb of interest poured into 60 mm Petri dishes (n = 3 per treatment group) and allowed to set at room temperature for 30 min. Individual colonies of ATCC 14028 STm were then picked from a freshly-streaked LB agar plate and stabbed directly into the center of the plate [22]. The plates were placed in a 37°C incubator and the diameters of the concentrically growing bacterial cultures were measured at 60-minute intervals.

### Bacterial agglutination assays

An aliquot of overnight liquid cultures of STm (100 μl) were mixed in equal ratio (v/v) with PBS containing a final concentration of 15 μg/ml of each IgA mAb of interest (n = 6 per treatment), and then placed into individual wells of a U-bottom 96 well plate. The plate was incubated at 37°C and visually monitored every 15 minutes for clumping of cells, as described [17].

### HeLa cell invasion assay

HeLa cells were obtained from the ATCC and maintained in Dulbecco’s Modified Eagle Media (DMEM) with 10% fetal bovine serum at 37°C and 5% CO_2_. Cells were seeded at 5 x 10^5^ cells/mL in 96-well plates and grown for 24 to 36 h to establish 70-90% confluency. Prior to STm infection assays, HeLa cells were washed three times with serum-free DMEM. Overnight cultures of AR05 and AR04 were subcultured in LB at 37°C with aeration and adjusted to an OD_600_ of ~0.7. Strains were mixed 1:1 and washed twice by centrifugation (6,000 x g for 2 minutes) and resuspended in PBS. Bacteria were then diluted 1:10 in Hank’s Balanced Salt Solution (HBSS, Wadsworth Center Media Core) and an aliquot was plated on LB agar supplemented with kanamycin (50 μg/mL) and X-gal (40 μg/mL) to compute bacterial input. For the invasion assay, bacterial mixtures were incubated with 15 ug/mL of either 2D6 IgA, PeA3 IgA, or Sal4 IgA mAb for 15 min at 37°C to minimize agglutination. Treated bacteria were applied to HeLa cell monolayers and centrifuged at 1,000 x g for 10 min (rotating the plate 180° at 5 min) to promote STm adherence to HeLa cell surfaces. The microtiter plates were then incubated for 90 minutes at 37°C. Cells were washed three times with HBSS and treated with gentamicin (40 μg/mL) to eliminate extracellular bacteria. Finally, cells were washed with HBSS lysed with 1% Triton X-100 (in Ca^2+^ and Mg^2+^-free PBS), serially diluted, plated on LB agar containing kanamycin (50μg/mL) and X-gal (5-bromo-4-chloro-3-indolyl-β-D-galactopyranoside) (40μg/mL) and incubated overnight at 37°C. The competitive index [(%strain A recovered/%strain B recovered)/(%strain A inoculated/%strain B inoculated)] was calculated for each treatment group.

### STm intragastric challenge

Overnight cultures of AR05 and AR04 were subcultured to an OD_600_ of 0.7, combined 1:1 (v/v) and resuspended in PBS. An aliquot was plated on LB agar containing kanamycin (50 μg/mL) and X-gal (40 μg/mL) at the start of the experiment to determine bacterial input (CFUs). Before gavage, bacteria (~4 x 10^7^ CFUs) were either incubated for 10 minutes with mAbs (30 μg/mL, unless stated otherwise), or mAbs were provided as a “chase” immediately before STm gavage (50 μg/mouse, unless stated otherwise). Twenty-four hours later, the mice were euthanized by CO_2_ asphyxiation followed by cervical dislocation. For each mouse, laparotomy was performed, and the small intestine was removed above the cecum. Peyer’s patches from each mouse were collected and placed in 1 mL sterile PBS on ice. Samples were then homogenized with a Bead Mill 4 Homogenizer (Fisher Scientific) three times for 30 seconds each. Homogenates were serially diluted, plated on LB agar containing kanamycin and X-Gal and incubated overnight at 37°C. Blue and white colonies were enumerated and the competitive indices (CI) were calculated as CI = [(% strain A recovered/% strain B recovered)/(% strain A inoculated/% strain B inoculated)]. Whole-plate dilutions (100 μl per plate) were required to observe enough colonies to calculate competitive indices. All samples that contained less than 30 CFUs (per 100 μl) were eliminated from the data set and considered “too few to count” (TFC). This pool of cells is more likely a representation of only a few bacteria that have successfully invaded and replicated within the lymphoid tissues, as Peyer’s patch entry by *Salmonella* has been shown the bottleneck for further dissemination during infection [16].

### Mouse model of STm systemic challenge

BALB/c female mice were administered 40 μg or 10 μg (unless otherwise indicated) of Sal4 IgA, Sal4 IgG1, or isotype control antibody in sterile PBS by intraperitoneal (i.p.) injection. An overnight culture of wildtype STm (ATCC 14028) was subcultured, washed in sterile PBS, and diluted to a final concentration of 5 x 10^4^ CFUs/mL. Twenty-four hours after passive immunization, mice were challenged with STm inoculum (1 x 10^4^ CFUs) by i.p. injection. An additional twenty-four hours later, mice were euthanized by CO_2_ inhalation followed by cervical dislocation and spleens and livers were collected, weighed, and homogenized in 1 mL sterile PBS as described above. Homogenates were serially diluted, plated on LB agar and incubated overnight at 37°C. Total CFUs were counted and computed for Log_10_ CFUs/gram (tissue).

### *In vitro* digestion assay

Sal4 IgA or Sal4 IgG mAbs were diluted to a final concentration of 0.1 mg/mL in simplified adult simulated gastric fluid (94 mM NaCl, 13 mM KCl; pH adjusted to 3.0 with 1M HCl) with or without pepsin (2000U/mL) on ice similarly as described [23–25]. Samples were incubated statically at 37°C and aliquots were taken after 10 minutes, 30 minutes, and 60 minutes of incubation and neutralized on ice to a pH of 7.0 to 7.4 using 1M NaOH. Following neutralization, all samples were analyzed for binding of purified STm LPS by ELISA as previously described. Concentrations of Sal4 IgA and Sal4 IgG were quantified by establishing a standard curve using SoftMax Pro 5.2.

## Results

### Oral administration of Sal4 IgA protects mice against invasive STm

To explore the benefit of passively administered Sal4 IgA on reducing the invasion of Peyer’s patch tissues by STm, we developed a competitive infection assay using two STm strains, AR05 and AR04 [19]. AR05 is a kanamycin resistant derivative of the type strain ATCC 14028 (**Table 1**) that expresses the O5 antigen (O5-Ag). AR04 is a derivative of AR05 with a *Tn10* insertion in the acetyl transferase gene (*oafA126::*Tn*10d-Tc*) that abolishes the bacterium’s ability to express the O5 Ag. Therefore, Sal4 IgA reacts with AR05 but not AR04 (**S1 Fig**) [14, 17]. In addition, AR04 constitutively expresses β-galactosidase. Thus, AR05 (“white”) is readily distinguished from AR04 (“blue”) on when colonies grown on LB/X-Gal agar (**S2 Fig**). The competitive index (CI) is simply the ratio of AR05 to AR04 in the inoculum compared to the ratio of AR05 to AR04 recovered from Peyer’s patch tissues (see Materials and Methods).

We employed the competition assay in a mouse model of STm intestinal invasion of Peyer’s patch tissues, as shown in **S2 Fig**. Adult BALB/c mice were challenged by gavage with a 1-to-1 mixture of AR05 and AR04 and 24 h later Peyer’s patches collected along the entire length of the small intestine. The tissues were pooled and homogenized and the resulting homogenates were serially diluted onto LB agar containing kanamycin and X-Gal. Preliminary studies determined that an inoculum of 4 x 10^7^ total CFUs (1:1 AR05 and AR04) resulted in the recovery 10^2^-10^4^ CFUs per mouse. The experiments also revealed AR05 was slightly more invasive than AR04, as evidenced by a CI of ~1.2 to 1.5.

Using this model, we confirmed that pre-treatment of the STm inoculum with Sal4 IgA resulted in a dose-dependent reduction in the number of AR05 recovered in Peyer’s patch tissues (**Fig 1**). The highest dose of Sal4 IgA tested (12 μg) resulted in >4 log_10_ reduction in AR05 invasion efficiency, as compared to controls. The absolute number of AR04 recovered from the same Peyer’s patch tissues was unaffected by Sal4 IgA. The relative impact of Sal4 IgA on Peyer’s patch invasion was more apparent when the recovery values were expressed as a CI (**Fig 1**). By this metric, the addition of as little as 0.4 μg of Sal4 IgA rendered AR05 at a competitive disadvantage compared to AR04 (**Fig 1**). The addition of greater amounts of Sal4 further reduced the CI with a maximal reduction occurring at concentrations above 1.2 μg Sal4 IgA. Invasion of Peyer’s patch tissues by AR05 and AR04 was unaffected by 2D6, an *anti-Vibrio cholerae* IgA mAb that served as the isotype control for these studies [15, 22].

**Figure 1.**
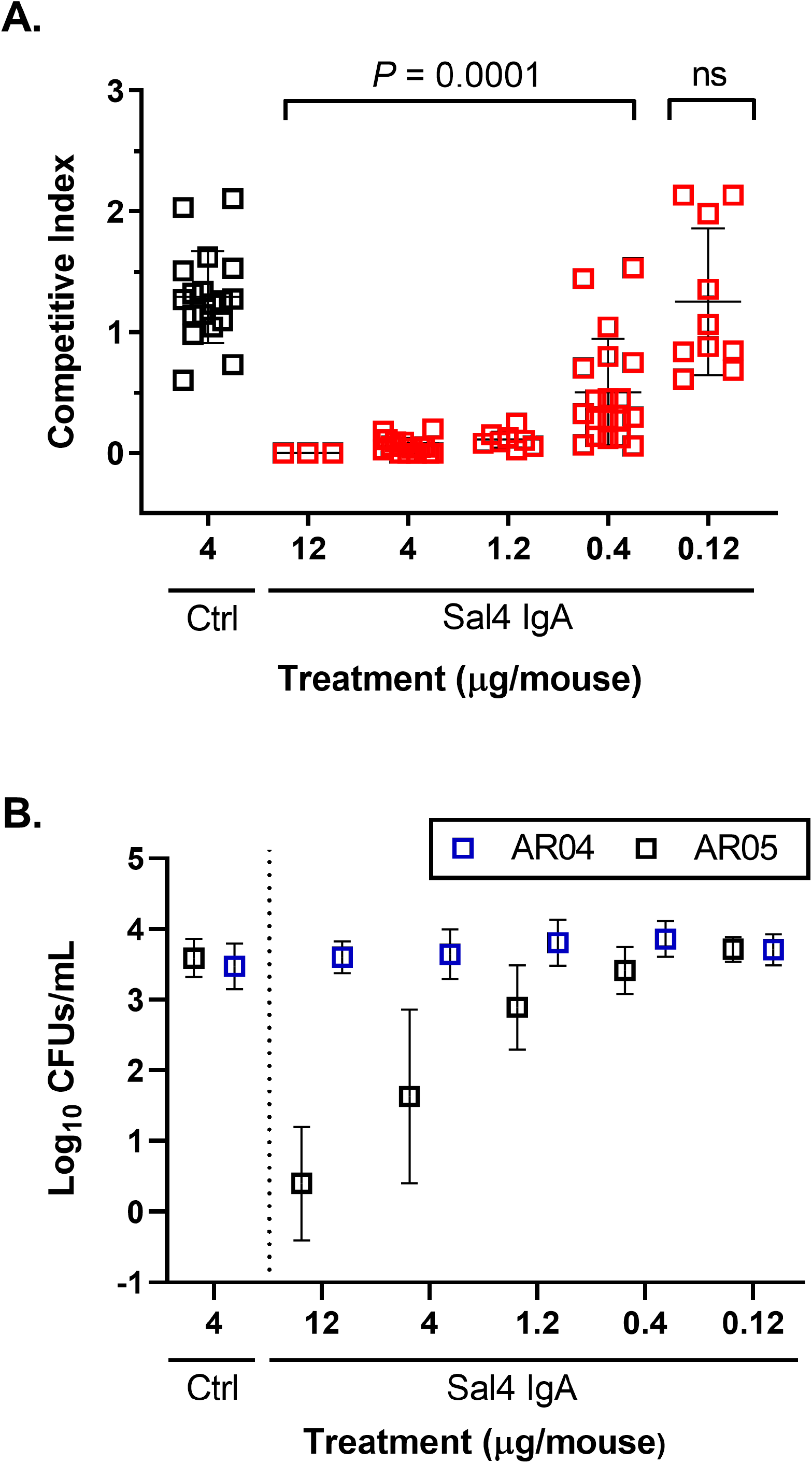
Orally administered Sal4 IgA blocks STm invasion into mouse Peyer’s patches. Adult BALB/c mice were challenged by gavage with a one-to-one mixture of STm O5 and O4 strains STm (4 x 10^7^ CFU total) co-administered with Sal4 IgA or an isotype control. Peyer’s patches were collected ~24 h later and assessed for bacterial loads. (A) Competitive indices and (B) total CFUs of wildtype and *ΔoafA* STm. Shown are the combined results of five independent experiments with at least 4 mice per group. Each symbol represents an individual mouse. Statistical significance evaluated for each concentration over the isotype control, as determined by Kruskal-Wallis test and Dunn’s post-hoc test.

However, Sal4 IgA’s activity was effectively lost when administered to mice in advance of STm challenge, possibly due to the deleterious effects of gastric pH or intestinal proteases like pepsin. For example, 36 μg of Sal4 IgA given to mice by gavage 20 min before STm challenge afforded no protection against Peyer’s patch uptake, as evidenced by an average CI of 1.2 (**S3 Fig**). In an effort to overcome this issue, the experiments were repeated with the addition of sodium bicarbonate (3% NaHCO_3_) or protease inhibitors. The co-administration of Sal4 IgA (50 μg) with either sodium bicarbonate or protease inhibitors resulted in a 40-50% reduction in AR05 uptake into the Peyer’s patches (**Figure 2**). While beyond the scope of the current study, these results underscore the necessity of identifying formulations capable of protecting orally administering antibodies from the gastric environment.

**Figure 2.**
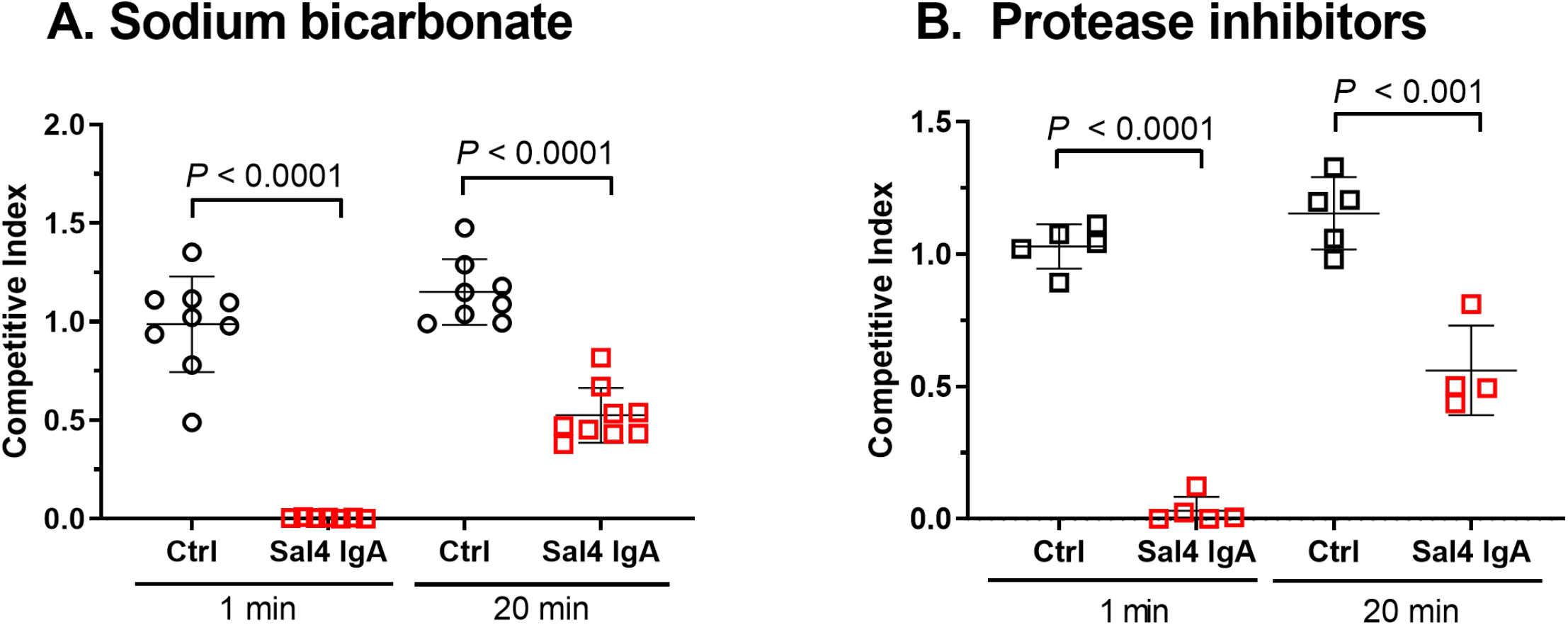
Sodium bicarbonate and protease inhibitors improve Sal4 IgA prophylactic activity. Adult BALB/c mice were gavaged with Sal4 IgA (50 μg) in (A) 3% NaHCO_3_ or (B) a protease inhibitor cocktail 20 min or 1 min before STm challenge. The mice were euthanized 24 h later and Peyer’s patches were assessed for bacterial loads, as a readout of bacterial invasion. Shown are the results of three independent experiments with at least 5 mice per group. Each symbol represents an individual mouse. Statistical significance compared to the isotype control at each time point, as determined by unpaired Student’s *t*-test.

### Benefit of SC on Sal4 IgA function in the mouse model

SC is reported to impart a number of biologically important activities upon SIgA in the context of the intestinal lumen, including protease resistance and mucus affinity [11, 26]. We therefore expected that Sal4 SIgA would be significantly more effective *in vivo* than equivalent amounts of dimeric Sal4 IgA lacking SC. To test this hypothesis, purified, dimeric Sal4 (dIgA) and dIgA complexed with human recombinant SC (SIgA) were compared side-by-side in the mouse model of invasive STm. Analysis of bacterial burdens in intestinal tissues collected 24 h after challenge revealed that Sal4 dIgA and SIgA were equally effective at limiting uptake of AR05 into Peyer’s patches (**Fig 3A**), indicating that the addition of SC did not enhance the protective capacity of Sal4 dIgA in this model.

**Figure 3.**
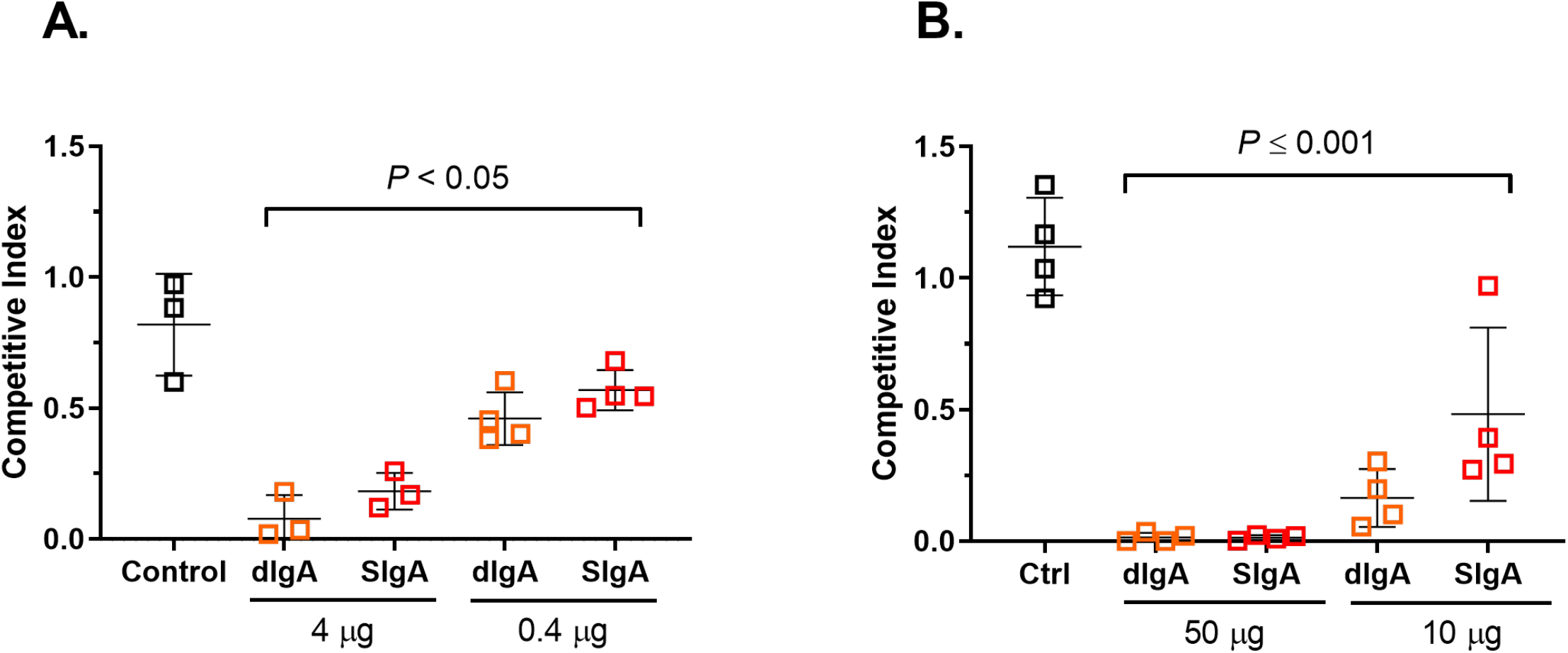
Comparison between Sal4 dimeric IgA (dIgA) and secretory IgA (SIgA) preparations in mouse model of STm infection. Adult BALB/c mice were challenged by gavage with a one-to-one mixture of STm O5 and O4 strains (4 x 10^7^ CFU total). Dimeric IgA (dIgA) or secretory IgA (SIgA) forms of Sal4 were (A) premixed with the bacterial inoculum or (B) administered ~1 min prior to STm challenge. The mice were euthanized 24 h later and Peyer’s patches were assessed for bacterial loads. Shown are the results of two independent experiments with 4 mice per group. Statistical significance evaluated for each concentration over the isotype control, as determined by one-way ANOVA and Tukey’s post-hoc test.

We postulated that the advantage of SC may only be apparent when antibody interacts with the intestinal environment in advance of bacterial challenge. We therefore repeated the experiments in which Sal4 dIgA and SIgA were given to mice by gavage immediately before STm challenge. Once again, however, Sal4 SIgA was no more effective than Sal4 dIgA at reducing invasion of AR05 into Peyer’s patch tissues. We conclude that, at least in this model of passive oral immunization, the potency of Sal4 IgA is not enhanced by the addition of SC (**Fig 3B**).

### Contribution of IgA avidity in inhibiting STm invasion of epithelial cells *in vitro* and Peyer’s patch tissues *in vivo*

While antibody avidity has been cited as an important parameter in controlling *Salmonella* infection in the gut following vaccination [27], its significance in the context passive oral immunization has not been examined. For instance, Sal4 IgA is the only O5-specific mAb that has been comprehensively characterized for biological activity *in vitro* and *in vivo* [12, 14, 15, 17, 19, 28]. We are unaware of experiments in the literature that have directly compared anti-STm IgA mAbs of differing avidities but with similar (or identical) epitope specificities.

We therefore sought to generate additional O5-specific mouse IgA mAbs and evaluate them for the ability to attenuate STm *in vitro* and *in vivo*. Toward this goal, groups of BALB/c mice were immunized with the an attenuated STm strain encoding a constitutively active PhoP mutant, as originally described [15]. B cell hybridomas were generated from Peyer’s patch lymphocytes from immunized mice and then screened by ELISA for IgA reactivity with STm strain 14028. Despite numerous attempts, we identified only one stable B cell hybridoma secreting an O5-specific IgA mAb, which we designated PeA3. PeA3 IgA bound STm LPS by ELISA, but with ~10-fold lower apparent avidity than Sal4 IgA (**S4 Fig**). In liquid culture, PeA3 agglutinated AR05, but not AR04, demonstrating its specificity for the O5 epitope (**S4 Fig**). PeA3 also bound the PhoP constitutive mutant strain by whole-cell ELISA, CS022, which was used as the immunogen (**S4 Fig**). In addition, PeA3 IgA was able to significantly impede STm flagella-based motility through soft agar, over a 6 h period (**S4 Fig**).

We next examined PeA3 IgA for the ability able to block *Salmonella* pathogenicity island 1 (SPI-1) type III secretion system (T3SS)-mediated entry of STm into HeLa cells [17]. A 1:1 mixture of AR05 and AR04 was treated with PeA3, Sal4, or the isotype control, 2D6, before being applied to HeLa cells with gentle centrifugation to bypass the need for motility in the invasion assay [17]. At the dose of antibody tested, PeA3 IgA treatment resulted in a significant reduction in AR05 invasion of HeLa cells, although its efficacy was slightly less than that of Sal4 IgA (**Fig 4**).

**Figure 4.**
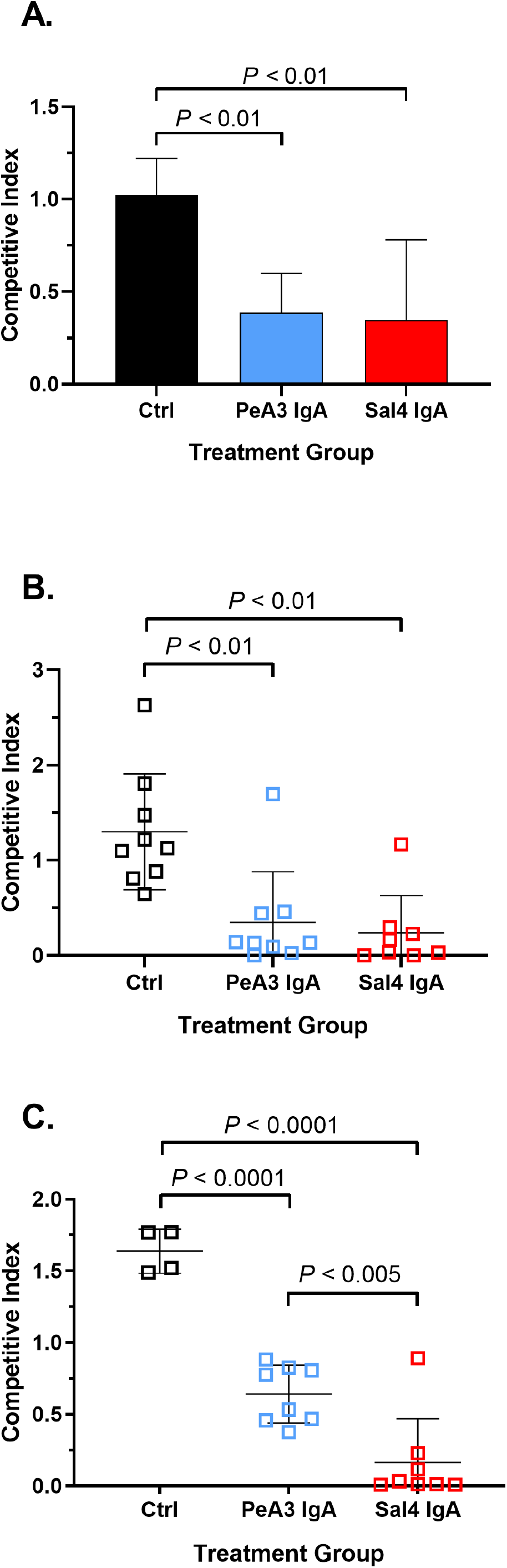
PeA3 IgA blocks wildtype STm invasion *in* and *in vivo*. (A) A competitive index of STm was treated with antibody and applied to HeLa cell monolayers to allow for invasion. Monolayers were treated with gentamicin to eliminate extracellular bacteria, then HeLa cells were lysed and enumerated for CFUs (*n* = 3 experiments done in triplicate). (B & C) Adult BALB/c mice were challenged by gavage with a one-to-one mixture of STm O5 and O4 strains (4 x 10^7^ CFU total) with (B) 30 μg/mL or (B) 10 μg/mL of indicated mAb. The mice were euthanized 24 h later and Peyer’s patches were assessed for bacterial loads. Shown are the combined results of two independent experiments with at least 4 mice per group. Statistical significance evaluated for each concentration over the isotype control, as determined by one-way ANOVA and either Dunnett’s (A & B) or Tukey’s (C) post-hoc tests.

To assess PeA3 IgA in intestinal immunity, it was compared to Sal4 IgA for the ability to block STm entry into mouse Peyer’s patch tissues. Groups of BALB/c mice were challenged with a 1:1 mixture of AR05 and AR04 supplemented with a high (12 μg) or low (4 μg) dose of PeA3 or Sal4 IgA. At the high dose, PeA3 and Sal4 IgA were equally effective in blocking bacterial uptake into Peyer’s patches, as evidenced by similar competitive indices (**Fig 4**). However, at the low dose, Sal4 IgA was superior to PeA3, as evidenced by mean CI values of 0.16 (± 0.30) versus 0.64 (± 0.20), respectively (**Fig 4**). Thus, the relative avidities of Sal4 and PeA3 for STm *in vitro* mirrors their efficacy *in vivo*.

### Potential of Sal4 IgG to function in passive immunization by the oral route

In clinical trials, ingestion of bovine milk- or colostrum-derived immunoglobulins consisting mainly of IgG from immunized dairy cows is sufficient to significantly reduce ETEC infection in adult volunteers [7, 29], indicating a role for IgG in passive oral immunizations. In fact, in a recent report, orally delivered anti-colonization factor antigen CFA/I IgG and SIgA human mAbs were equally effective at blocking ETEC infection in a mouse model [9].

To investigate the potential of orally administered IgG to prevent STm invasion of Peyer’s patch tissues, we engineered a Sal4 IgG chimeric antibody in which the V_H_ and V_L_ domains of the Sal4 IgA were grafted onto a human IgG1 framework (**S5 Fig**). The resulting Sal4 IgG1 was expressed in *Nicotiana benthamiana* using so-called RAMP technology [22]. *In vitro*, the chimeric IgG1 mAb had the same biological activity as Sal4 IgA, including reactivity with STm LPS by ELISA (**S5 Fig**). Sal4 IgG also promoted agglutination of STm in liquid culture, though slightly less effectively than Sal4 IgA (i.e. more antibody and longer incubation time was required to achieve macroagglutination). In a soft agar motility assay, Sal4 IgG limited bacterial spread over the course of the 6 h experiment, although slightly less effectively than Sal4 IgA. In the HeLa cell invasion assay, Sal4 IgG and Sal4 IgA were more or less equivalent in their abilities to block AR05 uptake (**Fig 5**). We therefore conclude that the Sal4 IgG1 molecule has expected the biological activities associated with Sal4 IgA, at least *in vitro*.

**Figure 5.**
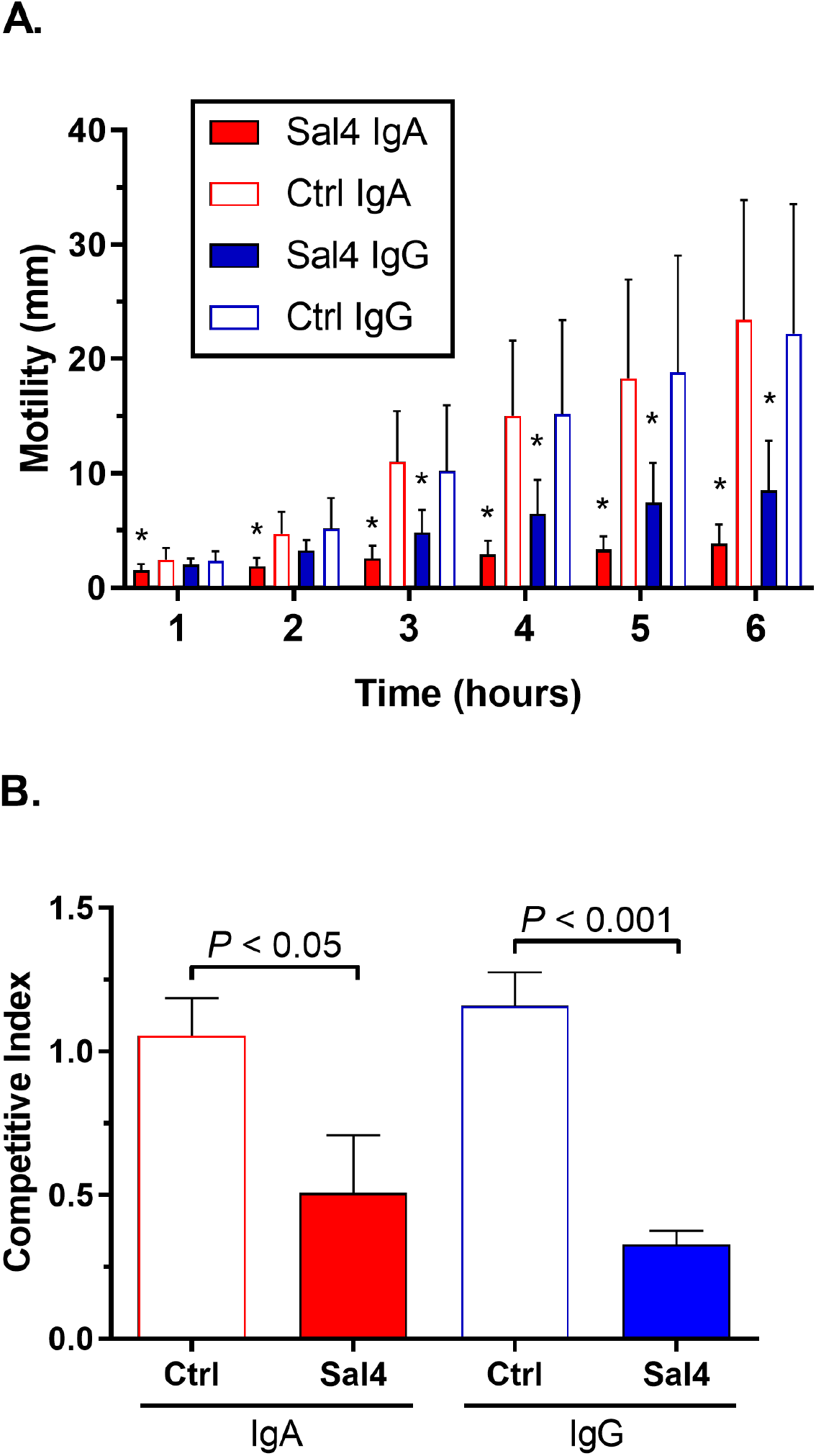
*In vitro* functionality of plant-based Sal4 IgG. (A) Effect of Sal4 IgG on STm motility in soft agar. STm was stab-inoculated into 0.3% LB agar containing 15 μg/mL of IgA control, Sal4 IgA, IgG control, or Sal4 IgG antibody. Plates were incubated at 37°C and the diameter of bacterial swimming was measured every hour for 6 hours (*n* = 3 experiments done in triplicate). (B) A one-to-one mixture of wildtype (AR05) and *oafA* mutant (AR04) STm (10^7^ CFUs) were incubated for 10 min with 15 μg/mL of IgA control, Sal4 IgA, IgG control, or Sal4 IgG antibody before being applied to HeLa cell monolayers (MOI 10), as described in the Materials and Methods. After 1 h, the HeLa cells were lysed and the CFUs were enumerated. Asterisks indicate significant reduction in wildtype STm motility (A) or invasion (B) over the respective isotype control, as determined by unpaired Student’s t-test (*n* = 2 experiments done in triplicate; *P* < 0.05).

We next investigated the potential of chimeric Sal4 IgG1 to passively immunize mice against STm infection in both systemic and oral challenge models. For the systemic challenge model, groups of BALB/c mice were administered Sal4 IgG1 or a chimeric IgG1 isotype control (PB10) by intraperitoneal injection and then challenged 24 h later with 10^4^ CFUs of wild type STm ATCC14028 by the same route. One day later, the mice were euthanized and CFUs in the spleens and livers were evaluated. As compared to the IgG1 control group, mice that received the high dose Sal4 IgG1 (40 μg) had 10 to 100-fold lower STm burden in the spleens and livers (**S6 Fig**). Bacterial numbers were also reduced in mice that received a low dose (10 μg) of Sal4 IgG1, although to a lesser extent than the high dose group of animals. These results demonstrate that passively administered Sal4 IgG1 results in dose-dependent reduction in STm systemic infection.

To examine Sal4 IgG1’s activity in the context of intestinal immunity to STm, groups of BALB/c mice were gavaged with a 1:1 mixture of AR05 and AR04 supplemented with 30 μg/mL Sal4 IgA or IgG1 or the relevant isotype controls. Invasion of Peyer’s patch tissues was measured 24 h later. As observed previously, Sal4 IgA reduced AR05 invasion into Peyer’s patch tissues by several orders of magnitude (CI value of 0.04 ± 0.02) (**Fig 6**). In contrast, Sal4 IgG1 had no effect on STm invasion, as evidenced by a CI value of 0.95 ± 0.13 (**Fig 6**). The failure of Sal4 IgG1 to function in these studies could not be overcome by increasing antibody dose (e.g., >750 μg) or repeated administration over a 12 h period (**S7 Fig**).

**Figure 6:**
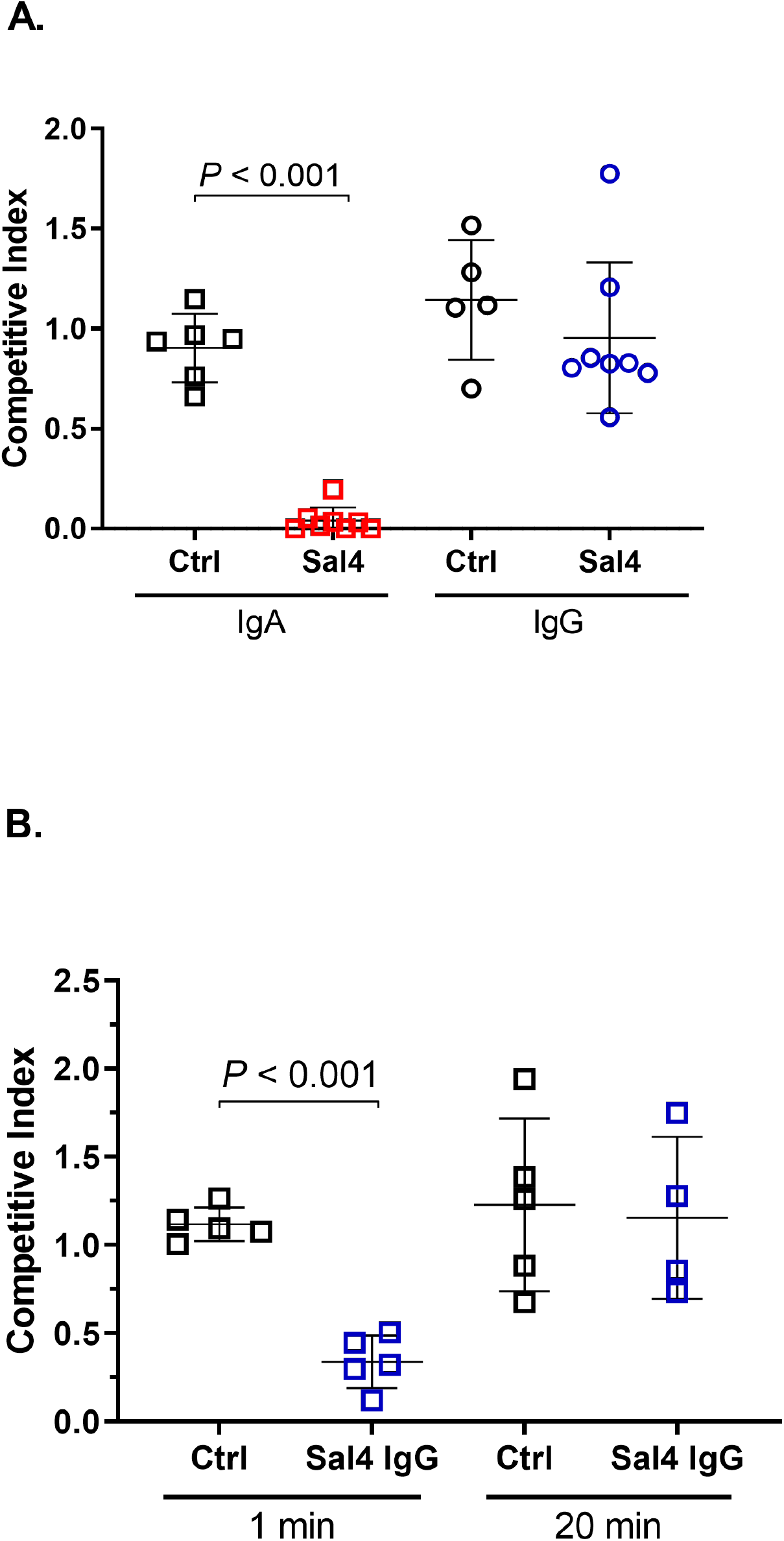
Sal4 IgG fails to protect against STm invasion following oral infection *in vivo but* can be partially rescued upon stabilization. BALB/c females were orally challenged with a competitive index of wildtype (AR05) and *oafA* mutant (AR05) STm (4 x 10^7^ CFUs) either (A) pre-incubated with 30 μg/mL of antibody or (B) administered antibody prior to STm challenge at the indicated time points. 24 hours (p.i.) mice were euthanized and Peyer’s patches were harvested and enumerated for CFUs and competitive indices. Asterisks indicate significant reduction in wildtype STm invasion over the control, as determined by Student’s t-test (*P* < 0.05).

We postulated that antibody stability in the gastric environment might account (at least in part) for the failure of Sal4 IgG1 to function in the oral passive immunization model [11, 25]. To address this experimentally, Sal4 IgA and IgG1 variants were incubated in an adult simplified Simulated Gastric Fluid (SGF), without and with pepsin, essentially as described [23, 24]. After 10, 30, and 60 min at 37°C, aliquots were removed and tested reactivity with STm LPS by ELISA. In the presence of SGF, Sal4 IgG1 levels declined steadily over a 30 min time period, whereas the IgA1 variant was relatively stable (**S1 Table**). Upon the addition of pepsin, however, IgG1 declined so precipitously that it was undetectable at 10 min. At the same time point (10 min), Sal4 IgA had declined to just ~10% of starting levels, but then remained detectable until 30 min. Collectively, these results confirmed the differential sensitivity of Sal4 IgG1 and IgA to the gastric environment.

The in vitro stability studies with Sal4 IgG prompted us to repeat the passive immunization studies with the addition of sodium bicarbonate plus protease inhibitors. Specifically, Sal4 IgG1 in sodium bicarbonate (3% NaHCO3) plus protease inhibitors administered to mice by gavage 1 min or 20 min prior STm challenge. Under these conditions, Sal4 IgG1 did in fact block STm invasion into Peyer’s patch tissues, but only when given immediately before STm challenge (**Fig 6**). Collectively, these results suggest that ineffectiveness of Sal4 IgG1, as compared to Sal4 IgA, is due to its instability in the gut environment.

## Discussion

In this study, we investigated the potential of orally administered mAbs to passively immunize mice against invasive Salmonella. The study was motivated by several factors. First is the rapid emergence of multi-drug resistance *Salmonella* infections, which constitute an increasing threat to public health in developing and even developed countries [30, 31]. Second, given the difficulty and extended timeline associated with the vaccine development, there is pressure from federal and private foundations to explore alternative strategies as a means of protecting at risk individuals from debilitating enteric infections. With the remarkable advances in recombinant mAb engineering and scale-up using mammalian cells, transgenic animals, plants and even seed-based [32] production platforms, the prospect of combatting diarrheal diseases through orally administered mAb cocktails is technically feasible and cost-effective.

We found that direct administration of Sal4 IgA to adult mice by gavage overcomes many of the impediments associated with the so-called “backpack” tumor model that was used previously to study STm-IgA interactions [33]. In the backpack model, antibody-secreting B cell hybridomas are implanted subcutaneously into mice, resulting in local tumor formation and the accumulation of antibodies at very high concentrations in serum and interstitial fluids, including the lamina propria [15]. Ultimately, hybridoma-derived, antigen-specific IgA is detected in intestinal secretions, presumably as a result of pIgR-mediated transcytosis [34, 35]. While this set-up resulted in physiologic delivery of Sal4 IgA into the intestinal lumen, the are several drawbacks with the model. First, depending on how well the hybridoma “takes”, the amount of Sal4 IgA in serum and intestinal secretions varies widely from mouse to mouse, thereby confounding the ability to perform dose-response studies. Second, because hybridoma-derived antibodies accumulate at potentially very high levels in serum (1-10 mg/ml) and interstitial fluids, it is not always possible to delineate whether observed protection is due to intestinal (secretory) or interstitial antibodies [34]. Finally, while it is assumed Sal4 IgA antibodies detected in the intestinal secretions in the backpack tumor mice are complexed with SC, the actual amount of Sal4 SIgA in the lumen has never been determined. Direct administration of Sal4 IgA of known molecular forms and at specific doses overcomes these concerns.

The other notable benefit of the challenge model employed here is that the primary readout is bacterial load (CFUs) in Peyer’s patch tissues, which are the primary point of entry for invasive *Salmonella* [36, 37]. Uptake into Peyer’s patches occurs through M cells and is dependent on the SPI-1 T3SS. Moreover, Peyer’s patch invasion occurs in the presence of a normal gut microbiota. This is in contrast to models of Salmonella-induced inflammation where infection occurs primarily in the cecum and colon and involves pre-treatment of mice with antibiotics like streptomycin to deplete the gut microbiota [38, 39]. For the purposes of this study, the challenge model granted us the ability to ask vital questions about IgA biology, such as the importance of SC in intestinal immunity.

Indeed, we found that the addition of SC did not augment Sal4 IgA activity *in vivo*, which is contrary to what we would have predicted based on the literature [10]. One possible explanation for why the benefits normally provided to IgA by SC were not observable in the context of the STm challenge model relates to the route of antibody delivery. Normally, dIgA is transported across the intestinal epithelium by the pIgR, which is preferentially expressed by enterocytes in intestinal crypts. Following transport, SIgA localizes to the mucus layer overlying the epithelial barrier where SC plays a central role in anchoring IgA within this microenvironment and protecting antibody from protease-mediated degradation [21, 40, 41]. It is unclear if the physiologic distribution of SIgA is recapitulated when antibody is administered by gavage. Our attempts to track, using immunohistochemistry, Sal4 SIgA in the small intestine following oral delivery have not been successful to date. Another possible explanation for why SC did not impart a benefit to Sal4 IgA is that the rate-limiting determinant for antibody-mediated protection in this model is dilution effects upon gavage, not protease sensitivity or mucus anchoring, where SC would be expected to play an important role.

The comparison between PeA3 and Sal4 IgA mAbs in the mouse model of invasive Salmonella serves as a demonstration of the importance of IgA avidity in protecting against invasive pathogens. By all accounts, Sal4 and PeA3 recognized the same or a very similar epitopes, but differed in their relative binding affinities for the O5 antigen by ~10 fold. The difference in binding activity correlated with differences in *in vivo* efficacy, at least when PeA3 and Sal4 IgA were given to mice at lower doses. At higher doses, Sal4 and PeA3 were equally effective at limiting STm uptake into Peyer’s patch tissues, underscoring that protection is the due to the interrelationship between IgA avidity and local antibody concentrations. As illustrated by Cothésy et al., even polyreactive SIgA is protective if present a sufficiently high concentrations [11]. From the standpoint of passive immunization, however, higher affinity/avidity antibodies are clearly advantageous since much lower doses would be required to achieve protection. Indeed, in the case of respiratory infections, the selection for higher affinity mAbs resulted in correspondingly higher neutralizing activities and in *vivo* potency [42].

It was disappointing (albeit not surprising) to discover that an IgG1 variant of Sal4 had a marginal capacity (and only when given with protease inhibitors and sodium bicarbonate) to passively immunize mice against intragastric Salmonella infection. Indeed, our results are consistent with other reports that demonstrate IgG1 instability in the gastric environment is a major contributor to its failure to function in the gut against Salmonella and other pathogens. It is likely that the heavily glycosylated nature of IgA provides an advantage upon direct delivery into the gut [43, 44] in terms of maintaining both direct antigen binding and crosslinking between multiple antigens [40], while the IgG mAb, with a lone pair of N-glycans on the Fc region [45], is outmatched. Other factors may also be at play. Sal4 IgG1, which is a monomer, likely differs from Sal4 IgA, which is a dimer, in its ability to promote bacterial agglutination. We cannot rule out the possibility that the nature of agglutination between IgG and IgA is quantitatively different considering that we did observe slight differences in the kinetics of microagglutination between to two antibody isotypes.

## Acknowledgements

We gratefully acknowledge Dr. Blaise Corthésy for providing dimeric Sal4 IgA and recombinant SC for this study. We thank Drs. Ozan Kumar, Sangeeta Joshi and David Volkin (University of Kansas) for assistance with simulated gastric fluids, and Dr. Charles Shoemaker (Tufts University) for rabbit intestinal fluids. We are grateful to Danielle Baranova and Dylan Ehrbar (Wadsworth Center) for providing control IgA antibodies and assistance with statistical analysis. We thank the Wadsworth Center’s Media and Tissue Culture Core facility for preparing media for mammalian and bacterial cultures. Finally, we extend our gratitude to Drs. Jeremy Blum and Omar Vandal (Bill and Melinda Gates Foundation) for their support of the program and for providing critical feedback and discussions throughout the study. This work was supported by grants to NJM from the National Institutes of Allergy and Infectious Diseases (AI119647) and the Bill and Melinda Gates Foundation (OPP1176017).

